# Conservation of antiviral systems across domains of life reveals novel immune mechanisms in humans

**DOI:** 10.1101/2022.12.12.520048

**Authors:** Jean Cury, Ernest Mordret, Veronica Hernandez Trejo, Florian Tesson, Gal Ofir, Enzo Z. Poirier, Aude Bernheim

**Affiliations:** Molecular Diversity of Microbes, INSERM U1284, Université Paris-Cité, Paris, France; Stem Cell Immunity Team, Institut Curie, PSL Research University, INSERM U932, Paris, France; Department of Molecular Biology, Max Planck Institute for Biology Tübingen, Tübingen, Germany

**Author notes:** These authors contributed equally (Jean Cury, Ernest Mordret).

## Abstract

Viral infection is a common threat to prokaryotic and eukaryotic life, which has resulted in the evolution of a myriad of antiviral systems. Some of these eukaryotic systems are thought to have evolved from prokaryotic antiphage proteins, with which they may display sequence and structural homology. Here, we show that homologs of recently discovered antiphage systems are widespread in eukaryotes. We demonstrate that such homologs can retain a function in immunity by unveiling that eukaryotic proteins of the anti-transposon piRNA pathway display domain homology with the antiphage system Mokosh. We further utilise this conservation to discover novel human antiviral genes related to the Eleos and Lamassu prokaryotic systems. We propose that comparative immunology across domains of life can be leveraged to discover immune genes in eukaryotes.

## Main

A common denominator of prokaryotic and eukaryotic life is the threat posed by viruses. In both domains, evolutionary pressures resulting from viral infections led to the selection of antiviral mechanisms. In chordates, including humans, the first line of defense against infection relies on the steady-state expression of restriction factors, as well as on the interferon-driven upregulation of a panel of antiviral effectors termed interferon-stimulated genes (ISGs) (*1*). Prokaryotes implement diverse antiviral systems to thwart bacteriophages (phages), including restriction-modification and CRISPR-Cas systems (*2*–*5*). Each system may be composed of a unique gene or of multiple genes whose products act in concert. Genes of a given system are closely located on the genome, allowing tight coregulation of gene expression and genetic exchange via horizontal gene transfer (*6*, *7*). Recently, more than 100 novel antiviral prokaryotic systems were uncovered (*2*–*7*). Among them, multiple bacterial antiviral systems were identified as the evolutionary origin of eukaryotic antiviral proteins, including CBASS (homolog of the eukaryotic cGAS/STING pathway), gasdermins, viperins or Avs (related to the NLR family) (*8*–*12*). In this work, we explore whether novel eukaryotic immune components can be identified by homology with prokaryotic defense systems.

### Eukaryotic genomes encode distant homologs of antiphage systems

We hypothesized that components of some of the recently-discovered antiphage systems may be conserved in eukaryotes, and thus could be identified through protein homology search. We used DefenseFinder, a pipeline developed to map known antiphage systems in prokaryotic genomes (*2*). This pipeline currently detects 132 types of antiphage systems, and relies on protein sequence homology to detect a system’s individual components. Each component is searched using a protein profile allowing for low homology detection (hidden Markov model, HMM). Because most antiphage systems are organized in operons, DefenseFinder also takes into account genomic proximity on the chromosome (colocalization) of genes from a given system. Our analysis thus relies on two complementary criteria: 1) for single-gene systems, high sequence homology with a protein profile; 2) for multi-gene systems, a more relaxed sequence homology of each component coupled with the constraint of genes’ colocalization. These two criteria are expected to limit the potential detection of antiphage homologs in eukaryotes as 1) protein homology may be limited and 2) coregulation of eukaryotic gene expression does not canonically rely on genomic colocalization. Nonetheless, genes of eukaryotic homologs of antiphage systems can be genomically colocalized, as recently observed for viperin and its kinase partner in metazoans, including humans (*9*, *13*).

DefenseFinder analysis of a database of 4,616 genomes spanning the kingdom of eukaryotes detected 14,217 hits, which were filtered to remove bacterial genomic hits (Table S1 and Material and Methods). This approach was successful in detecting hits of eukaryotic immune proteins with known antiphage homologs, such as viperin (Fig. 1A, 1,164 homologs). Viperin’s antiphage activity relies on the generation of modified nucleotides that act as chain terminators of viral replication. In mammals, viperin is an ISG that stops infection by a similar mechanism (*9*, *14*). We also identified 34 instances of homology with CBASS, which is the prokaryotic homolog of the cGAS/STING pathway, a key antiviral immune pathway in animals (Fig. 1B). cGAS detects viral DNA and signals through STING to induce an antiviral response (*1*, *15*). In animals, STING is considered to be the canonical partner of cGAS. There is more diversity in bacteria, in which cGAS proteins can be associated with other effectors such as phospholipases (*8*). Our search documents gene colocalization between cGAS and phospholipases in arthropods, suggesting that other versions of the CBASS antiphage system may be present in eukaryotes (Table S1).

**Fig. 1.**
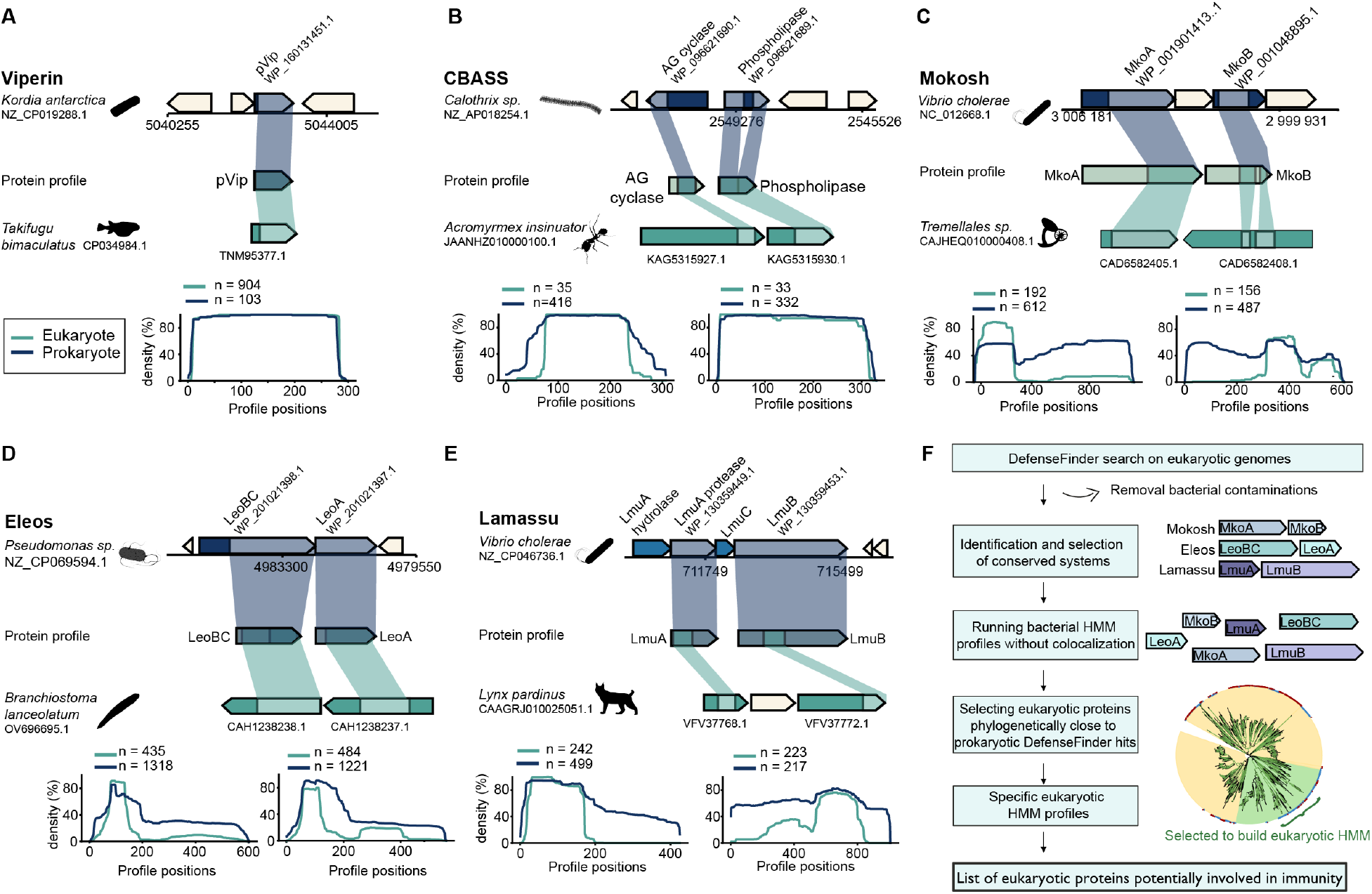
Distant homologs of antiphage systems can be detected in eukaryotes. (**A-E**) Representation of DefenseFinder hits for the antiphage systems viperin (A), CBASS (B), Mokosh (C), Eleos (D) and Lamassu (E) in their genomic context. Prokaryotic and eukaryotic proteins are colored in blue and green, respectively, and labeled with their NCBI identification number. Homology between eukaryotes and prokaryotes are highlighted by colored shades, mapping on a given HMM protein profile. Bottom plots quantify the average coverage of the HMM profile for all eukaryotic (green) and prokaryotic (blue) hits. Total number of unique hits are indicated. (**F**) Bioinformatics pipeline used in this study.

Because we detected known eukaryotic homologs of antiphage pathways, we then assessed if a similar approach could be leveraged to unravel additional conserved systems. We identified numerous homologs of the two-gene systems Mokosh (MkoA, MkoB), Eleos (LeoA, LeoBC) and Lamassu (LmuA, LmuB). 178 homologs of the Mokosh system were detected in 165 eukaryotic genomes (Fig. 1C). The antiphage system termed Eleos in this work, after the Greek goddess of compassion, corresponds to the combination of LeoA and LeoBC, which was previously described as an antiphage system with dynamin-like domains (*3*, *16*). 415 homologs of Eleos were identified in 317 eukaryotic genomes (Fig. 1D). Finally, 624 homologs of Lamassu were detected, notably in rotifers, arthropods and chordates (88% of the hits, Fig. 1E and S7) (*3*, *6*). These data suggest that unknown homologs of bacterial immune systems can be detected in eukaryotic genomes by using protein homology searches that are calibrated on bacterial proteins.

This initial analysis identified distant homologs of antiphage systems in eukaryotes. It however relies on a criterion of genomic colocalization, which prevents the detection of homologous multi-gene systems in which all components exist in a eukaryotic genome, but would not be genomically colocalized. Among the full systems detected by DefenseFinder in eukaryotic genomes, we focused on Mokosh, Eleos and Lamassu, and scanned our database of genomes for their individual components (Fig. 1F). The number of eukaryotic hits obtained with this analysis varied drastically between profiles, ranging from 1,080 for MkoB to 1,095,192 for MkoA (fig. S1-2 and Table S2). Since protein domains can typically be shared between immune and non-immune proteins (*17*), we expect the resulting hits to be a mix of immune and non-immune proteins. We hypothesized that eukaryotic proteins with immune roles would be phylogenetically closer to prokaryotic proteins with immune functions – defined, here, as DefenseFinder hits in prokaryotic genomes. We devised a procedure (see Material and Methods) to create a set of HMM profiles from groups of eukaryotic proteins, based on their phylogenetic proximity to prokaryotic immune proteins (fig. S1). We used these new HMM profiles to look for eukaryotic proteins potentially involved in immunity. These new profiles detected various numbers of hit, from 1,270 (MkoB) to 68,655 (LmuA) proteins in the eukaryotic genomes. With respect to our initial analysis relying on bacterial HMM profiles, our query using refined eukaryotic profiles is expected to retrieve a more specific set of hits, as it putatively encompasses the phylogenetic relationship to prokaryotic immune proteins. It is also expected to be more sensitive, as the profile should capture eukaryotic sequence diversity. As such, only 17.4% of hits are common between the two methods (fig. S3). Hits uncovered using eukaryotic profiles are described in the following sections (Tables S3, S4, S5), with a focus on humans (Table S6).

### Domains with structural homology to the antiphage Mokosh system participate in the anti-transposon piRNA pathway of the animal germline

As a proof-of-concept, we first determined if our comparative immunology approach allows to identify known human defense genes. Mokosh is an antiphage system composed of proteins with at least two domains: an RNA helicase (encoded by MkoA) and a phospholipase D domain (PLD, present in MkoB) (Fig. 1C and 2C) (*3*). Our analysis revealed a single human hit of MkoB: the mitochondrial cardiolipin hydrolase PLD6 (only identified using the eukaryotic HMM profile, fig. S3). This protein is involved in the piRNA pathway, a key defense mechanism of the animal germline (Fig. 2A,C, fig. S5A,B and Table S3) (*18*). piRNAs prevent the expression of transposable elements, which are genomic sequences originating, in part, from ancient events of retrovirus integration (fig. S5B). PLD6 is a nuclease responsible for the degradation of piRNA precursors, that are ultimately converted into mature piRNAs to target transposable elements (fig. S5B). It does so by interacting with the RNA helicase MOV10L1, which unwinds piRNA precursors (*18*–*20*). Our analysis shows that these two proteins, MOV10L1 and PLD6, share domains with the prokaryotic proteins MkoA and MkoB, respectively (Fig. 2A-C). To explore prokaryote-eukaryote structural conservation, we aligned the conserved domains of human PLD6 and MOV10L1 with a predicted Mokosh protein from *E. coli*, in which MkoA and MkoB are fused in a single protein (Fig. 2D). The PLD domain of PLD6 shows homology to *E. coli* Mokosh (template modeling score, TM-score of 0.63), as does the RNA helicase domain of MOV10L1 (TM-score of 0.74) (Fig. 2D,E). We expanded structural comparisons to other animals, in which the piRNA pathway relies on PLD6 and MOV10L1, and similarly showed that domains of both proteins have clear structural homologies with prokaryotic MkoB and MkoA (fig. S5D). While the RNA helicase and PLD domains are conserved in MOV10L1 and PLD6, respectively, the proteins also include additional domains that are not present in bacterial Mokosh systems (fig. S5D). This data suggests that the antiphage system Mokosh may have been the origin of these key proteins of the piRNA pathway, highlighting another case of a possible bacterial origin of a eukaryotic protective mechanism.

**Fig. 2.**
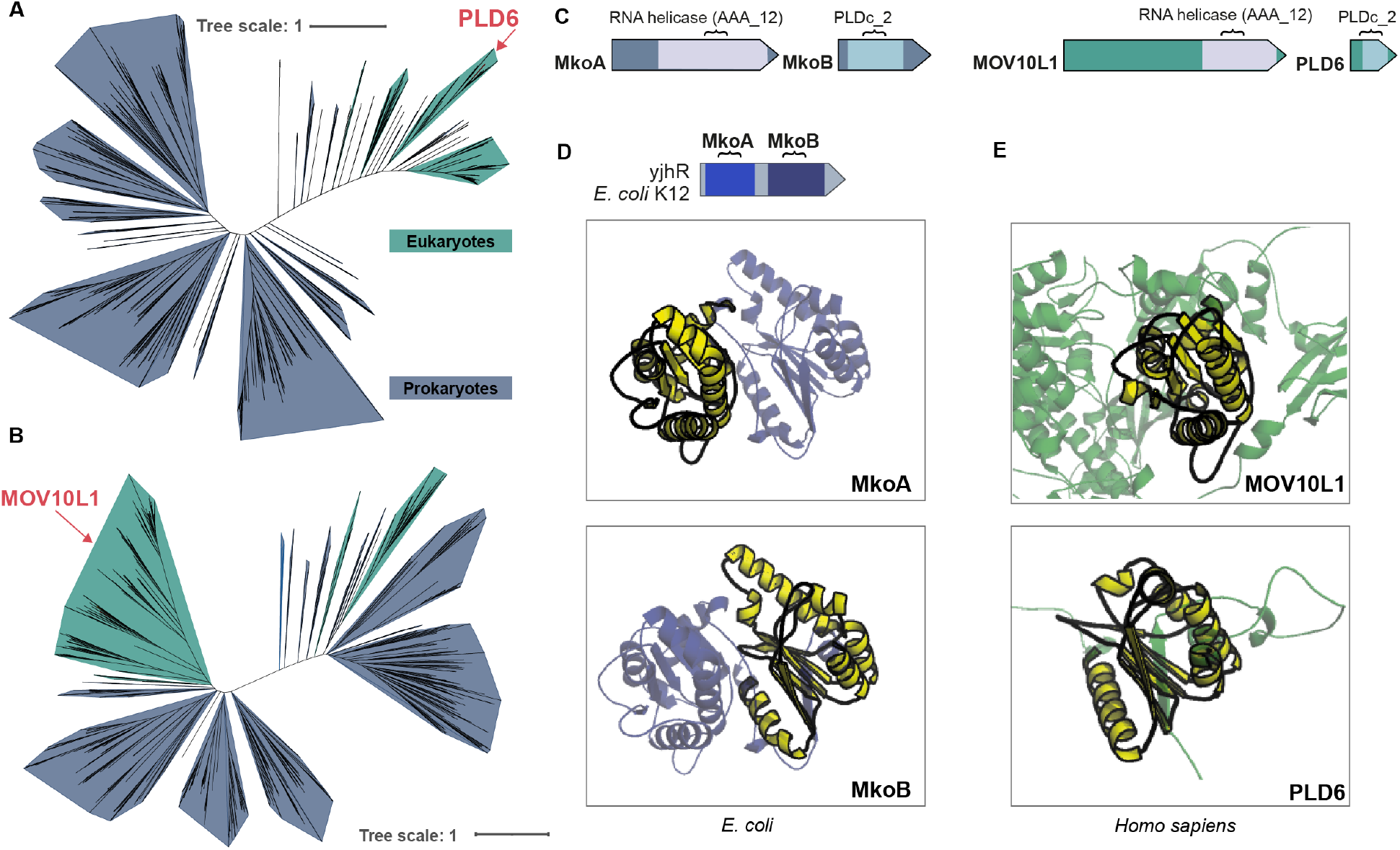
The protective piRNA pathway of the animal germline involves domains shared with the antiphage system Mokosh. (**A-B**) Phylogenetic analysis combining prokaryotic MkoA (A) and MkoB (B) with their eukaryotic hits (see Material and Methods). Branches are colored according to the kingdom (blue for prokaryotes, green for eukaryotes). Human hits of interest are highlighted in red. (**C**) Conservation of the RNA helicase and PLD-like nuclease domains of prokaryotic antiphage MkoA and MkoB (blue) in the human homologs MOV10L1 and PLD6 (green). (**D**) Structural comparison of a Mokosh from *E. coli* K12, in which MkoA and MkoB are fused (yjhR protein, depicted in blue), with the helicase domain of human MOV10L1 (top) or the PLD domain of human PLD6 (bottom). Optimal local alignment between the two structures (as determined by foldseek) is represented in yellow. (**E**) Structural comparisons of human MOV10L1 (top) and PLD6 (bottom) with the yjhR protein of *E. coli*. For MOV10L1 (respectively PLD6) homologs, the yellow domain corresponds to the protein’s optimal alignment to the MkoA (respectively MkoB) domain of yjhR. Part of the structures are omitted and shown in Fig S5. Structures were predicted using AlphaFold. PDB identifiers and TM-scores can be found in Table S7.

Note that our search uncovered 15 additional hits of MkoA, 27% of which have been described to be involved in antiviral immunity (Table S6). The ISG helicase with zinc finger domain 2 (HELZ2) is antiviral against dengue virus by regulating lipid metabolism through its interaction with the aryl hydrocarbon receptor (*21*). RNA helicase zinc finger NFX1-type containing 1 (ZNFX1) is a dsRNA sensor that triggers a type-I interferon response upon RNA virus infection (*22*). Altogether, this data provides a proof-of-principle that comparative immunology across domains of life is a fruitful method to identify defense genes in eukaryotes.

### GIMAPs, human proteins related to antiphage Eleos, are antiviral

We then asked if our approach could be leveraged to identify novel antiviral factors. Eleos is a two gene-system (LeoA and LeoBC) coding for proteins with dynamin-like domains, from the P-loop NTPase superfamily (*3*). The search for human homologs of LeoBC identified two protein families: EH domain-containing proteins (EHDs) and GTPases immunity-associated proteins (GIMAPs) (Fig. 3A,B and Table S6). We focused on GIMAPS, a group of 8 human paralogs, as they are transcriptionally induced by interferon, and have a documented role in T lymphocyte survival as well as in resistance to infection by the parasite *Toxoplasma gondii* (*23*–*25*). We hypothesized that GIMAPs could also be involved in antiviral immunity. Structural comparison between a LeoBC of *Bacillus sp*. HY001, and GIMAP6 revealed moderate structural similarities of the P-loop NTPase domain (predicted structures, TM-score of 0.45, Fig. 3C). Expression of human GIMAP5 and 6 was induced in human embryonic kidney 293T cells by plasmid transfection, and cells were infected with GFP-encoding herpes simplex virus 1 (HSV-1), a DNA virus. GIMAP expression, as well as HSV-1 infection, were monitored at the single cell level using intracellular immunostaining and flow cytometry (fig. S6A-C). Cells expressing GIMAP5 and 6 display 1.4- and 2-fold reduction in infection by herpes simplex virus 1, respectively (Fig. 3D). Viral replication measured by the intensity of virus-encoded GFP, was reduced by up to 40% (fig. S6D). To interrogate if GIMAP antiviral activity is herpes-specific, GIMAP-expressing cells were infected with Sindbis virus (SINV), an RNA virus. Viral infection and replication, measured by the expression of SINV-encoded GFP, were reduced by up to 2 folds upon expression of GIMAP5 and 6 (Fig. 3E and fig. S6E). We concluded that human proteins with domains similar to prokaryotic Eleos are ISGs that exert an antiviral activity against DNA and RNA viruses.

**Fig. 3.**
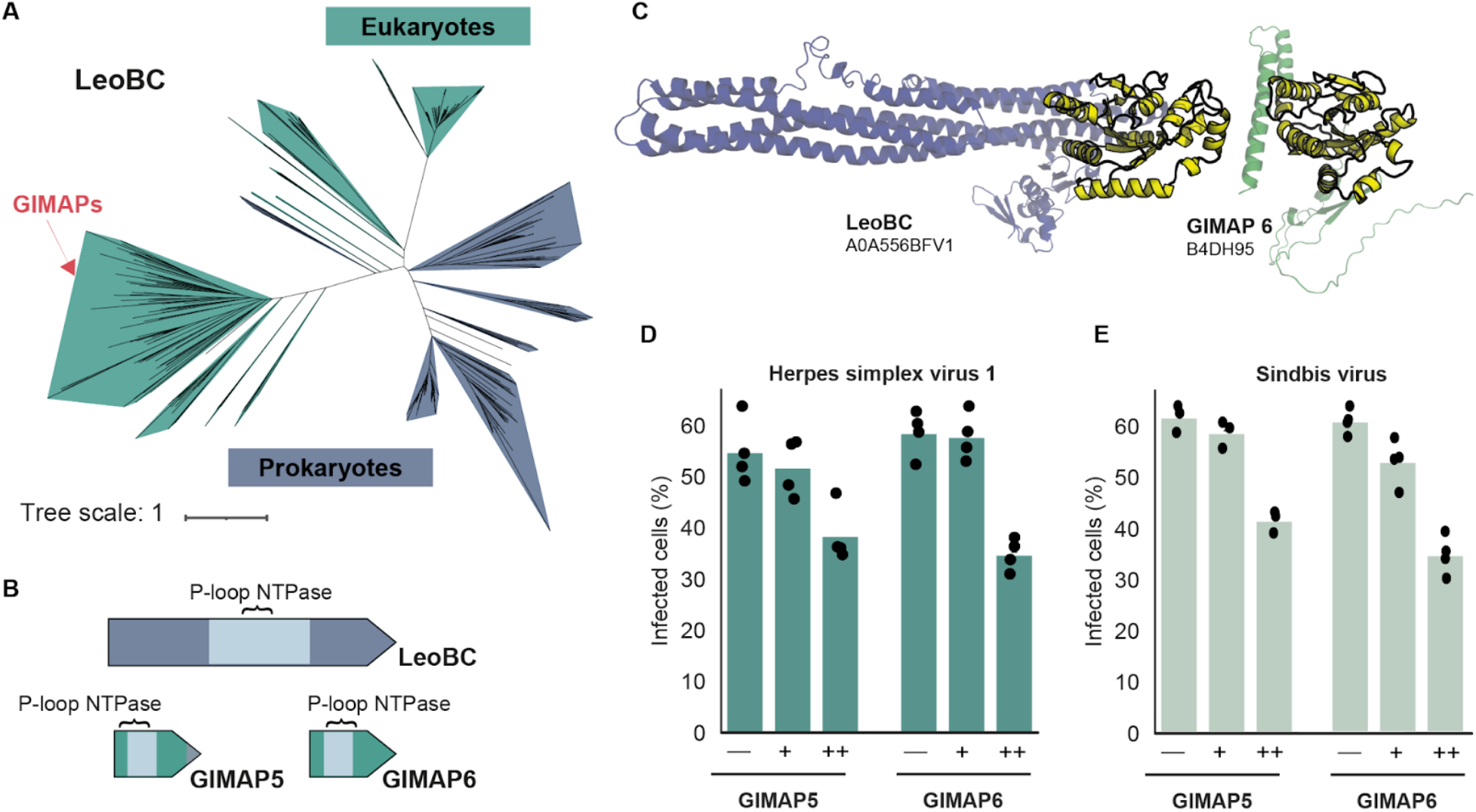
GIMAPs, proteins related to the antiphage Eleos system, are antiviral. (**A**) Phylogenetic analysis combining prokaryotic LeoBC with their eukaryotic hits. Branches are colored according to the kingdom (blue for prokaryotes, green for eukaryotes). Positions of human GIMAPs are indicated in red. (**B**) Conservation of the P-loop NTPase domain of prokaryotic antiphage LeoBC (blue) in human GIMAP5 and 6 (green). (**C**) Structural comparison of prokaryotic LeoBC with human GIMAP6. TM-score = 0.4. Structures were predicted using AlphaFold (Table S7). (**D,E**) 293T cells were transfected with plasmids encoding for GIMAP5 or GIMAP6 and infected with HSV-1 (D) or SINV (E) coding for GFP. Expression of GIMAPs and GFP was quantified by flow cytometry at 48h (HSV-1) or 24h (SINV) post-infection. GIMAP–, no expression; GIMAP+, mild expression; GIMAP++, high expression as described in fig. S6. Two-sided independent t-test comparing GIMAP– and GIMAP++, (D) p = 0.008 (GIMAP5) and p = 0.001 (GIMAP6), (E) p = 0.00005 (GIMAP5) and p = 0.0001 (GIMAP6).

### Distant human homologs of antiphage Lamassu are antiviral

Finally, we explored if novel human genes with no known links to immunity could be identified using comparative immunology across domains of life. One unexpected aspect of our analysis is the detection, in eukaryotic genomes, of gene colocalization for hits of the different components of a same system. Lamassu is composed of LmuB, which contains a SMC domain predicted to bind DNA, and LmuA, a predicted effector that can be a trypsin protease (Fig. 4A) (*3*). 35% of chordates genomes encode LmuA and LmuB colocalized (fig. S7). A protein fusion of trypsin domain (LmuA) and coil-coiled domain (LmuB) can be detected in the *Cyprinidae* family of fish (Fig. 4B and fig. S8A,B). The existence of fusion proteins in fish, as well as the conservation of genomic colocalization, suggest that the two proteins participate in the same pathway also in eukaryotes. In humans, out of 109 hits to LmuA and 28 proteins to LmuB, one genomic locus encodes both LmuA and LmuB colocalized.

**Fig. 4.**
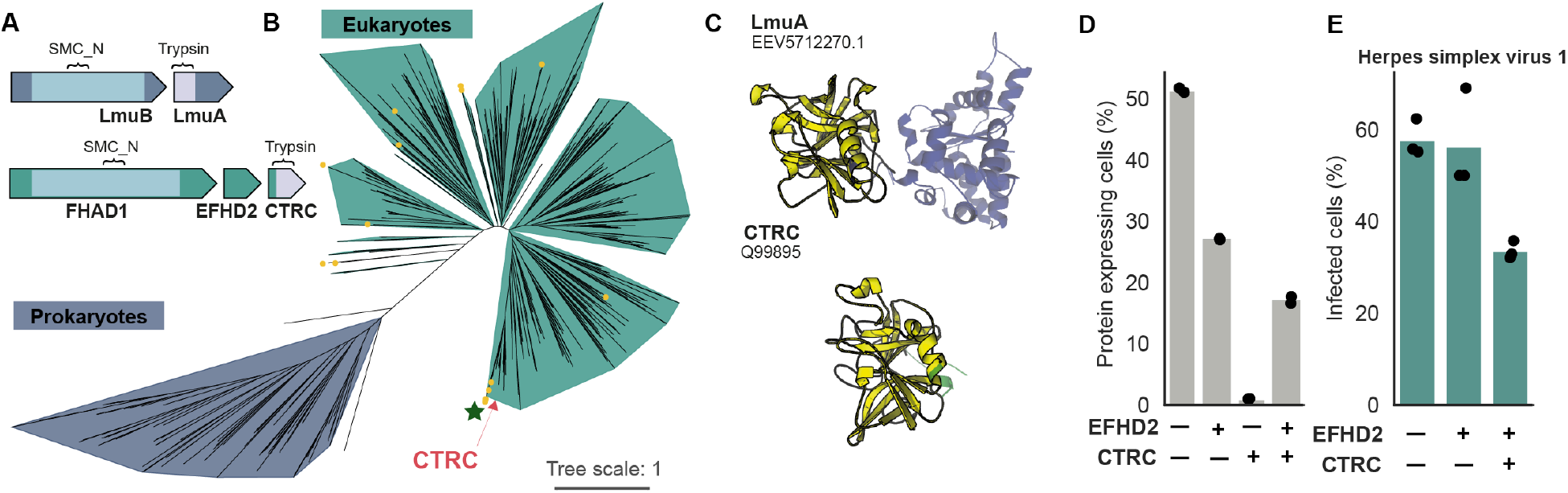
CTRC, a structural human homolog of the antiphage protein LmuA, is antiviral. (**A**) Conservation of the trypsin and SMC_N domains of prokaryotic antiphage LmuA and B (blue) in human CTRC and FHAD1 (green). (**B**) Phylogenetic analysis combining prokaryotic LmuA with its eukaryotic hits (see Material and Methods). Branches are colored according to the kingdom (blue for prokaryotes, green for eukaryotes). Human CTRC is indicated in red. Fusion proteins hit by both eukaryotic profiles are indicated by yellow dots. Fusions found in the *Cyprinidae* family are flagged with a green star. (**C**) Structural comparison of LmuA from a Lamassu of *E. coli* with human CTRC, TM-score = 0.69. Structures were computed using AlphaFold (Table S7). (**D,E**) 293T cells were transfected with plasmids encoding for EFHD2 or CTRC and infected with HSV-1. Expression of CTRC and EFHD2 (D) and virus-encoded GFP (E) was quantified by flow cytometry at 48h post infection. Two-sided independent t-test comparing EFHD2–CTRC– to EFHD2+CTRC+, (E) p = 0.0007.

Chymotrypsin-C (CTRC) protease is detected as a hit in our LmuA search, and the human *CTRC* gene is located next to *Forkhead-associated phosphopeptide binding domain 1 (FHAD1)*, whose protein product is a hit in our LmuB analysis (Fig. 4A-C and fig. S8B). *FHAD1* and *CTRC* genes are separated by *EF-hand domain-containing protein D2* (*EFHD2*, Fig. 4A). This protein performs pleiotropic functions, including regulation of immune activation in myeloid cells (*26*). CTRC is a protease secreted by pancreatic exocrine cells (*27*). It is additionally expressed, as is EFHD2, by macrophages and dendritic cells, responsible for immune surveillance and priming of the adaptive immune response (fig. S9). Intracellular levels of CTRC and EFHD2 were analyzed by immunostaining and flow cytometry in 293T cells transfected concomitantly with two plasmids encoding both proteins. As the transfection efficiency does not reach 100%, each cell will receive either no plasmid, only EFHD2 or CTRC plasmid, or both plasmids. Because CTRC is canonically secreted, one should not expect to detect intracellular accumulation of the protein in CTRC-transfected cells. Indeed, cells containing only CTRC protein represent 1% of total transfected cells (Fig. 4D and fig. S10A). However, cells expressing CTRC can be detected when coexpressing EFHD2 (18% of total transfected cells, Fig. 4D). This suggests that EFHD2 affects the secretion of CTRC and triggers its intracellular retention. 293T cells expressing CTRC+EFHD2, but not EFHD2 alone, display a two-fold reduction in infection with HSV-1 (Fig. 4E and fig. S10B). No clear antiviral effect was detected against SINV (fig. S10C,D). Addition of FHAD1 did not improve the antiviral capacity of EFHD2-CTRC in this experimental system, which could be related to the low expression of the protein compared to CTRC and EFHD2 (fig. S10E,F). Thus, the human protein CTRC, with distant homology to antiphage Lamassu, is a restriction factor that displays antiviral activity against a herpesvirus.

## Discussion

In this work, we unravel the conservation of recently discovered bacterial antiphage systems in eukaryotes, and utilize it to discover novel immune proteins in humans. These data – together with previously documented conservation events between prokaryotes and eukaryotes – show that proteins related to defense mechanisms in bacteria are involved at multiple levels of eukaryotic immune pathways, including pathogen detection (cGAS/STING), signal transduction (TIR domains), activity against transposons (PLD6/MOV10L1) and antiviral effectors (GIMAPs, EFHD2-CTRC) (*8*, *12*, *18*, *28*). Bacterial antiphage systems could thus have served as providers of building blocks for the evolution of immunity in eukaryotes. Importantly, the fact that certain hits in our search display antiviral activity does not imply that all hits in our search will have a role in immunity. Instead, a significant proportion may have been co-opted for unrelated cellular functions, or may present a dual role within and outside of immunity. Evolutionary and mechanistic studies will help delineate each protein’s contribution to eukaryotic immune pathways. We did not perform an exhaustive analysis of antiphage homologs, but rather chose to focus on specific systems. It is thus likely that additional homologs of antiphage systems will be identified.

Our work shows that three antiphage systems share domains with eukaryotic proteins participating in immunity. MOV10L1 and PLD6, reminiscent of Mokosh, play a key role in the piRNA pathway which protects the animal germline against transposable elements. Human proteins displaying domain conservation with Eleos (GIMAP5-6) and Lamassu (CTRC-EFHD2) can shield cells from HSV-1 infection. Interestingly, GIMAPs and CTRC-EFHD2 can block herpesviruses, which is part of the *Duplodnaviridae*, the realm of virus that includes bacteriophages (*29*). Connexions between phages and herpesviruses were indeed documented by a recent work describing the antiphage system Avs, that can be triggered *in vitro* by human herpesvirus-8 proteins (*11*).

Our data also documents a diversification of the proteins’ protective roles, which can be rewired to address other eukaryotic threats. The Mokosh system could have been co-opted to block transposable elements integrated in the genome. Similarly, GIMAP5 and 6 can also target RNA viruses, a broad antiviral tropism also documented for other members of the dynamin superfamily such as the ISGs MxA and MxB (*30*). There is thus a certain level of modularity in the evolutionary co-optation of antiphage systems, which may participate in various branches of eukaryotic immunity, including responses against bacteria or fungi. In addition to unraveling striking prokaryote-eukaryote conservation, our work proposes a comparative immunology approach to discover mechanisms of defense across domains of life, including phyla like plants or protists. We anticipate that the refinement of our method, as well as the expansion of knowledge on antiphage systems, will provide a more comprehensive understanding of prokaryotes’ contribution to eukaryotic immunity and lead to the identification of novel immune pathways across eukaryotes.

## Materials and Methods

### Database of eukaryotic and prokaryotic genomes

Eukaryotic genomes were downloaded from genbank in January 2022. Isoforms were removed from the protein fasta files, when possible, by keeping the longest one. Pseudogenes were removed as well, when annotated as such. CDS were renamed to take into account the relative position to easily assess whether two proteins are co-localized in the genome. In total, 4,616 eukaryotic genomes representing 2407 species were screened. 22,920 prokaryotic complete genomes were downloaded from Refseq in July 2022. List of genomes are available at https://github.com/mdmparis/cury_mordret_et_al_supplementary_data.

### Identification of homologous antiphage systems in eukaryotic genomes

DefenseFinder v1.0.9 with models v1.2.2 was run with default parameters on the custom database of 4,616 eukaryotic genomes, generating 14,701 candidate systems composed of a total of 17,932 associated genes. The output hits of DefenseFinder was examined to remove potential bacterial genomic contaminations. Each candidate gene was searched against the UniProt90 (https://www.uniprot.org) database using Diamond (v2.0.9 -b 10 -c 1 --max-target-seqs 5 --outfmt 6) (*31*), retrieving the 5 best hits per query. We used the uniprot id-mapping API to fetch the common ncbi taxon id of each of the UniProt90 families. This step fetched a taxon id for 78% of the hits. For the remaining 22% of the hits whose protein’s id did not match any UniProt90 family, we fetched their taxonomy by mapping the id to UniProtKB (12%) or to UniParc (10%). Each candidate protein was deemed a contamination if at least one of the 5 best hits on the UniProt90/UniProtKB/UniParc database came from a prokaryote, and its associated systems were subsequently removed from the pipeline. This approach led us to remove a total of 746 (4.2%) genes and 484 systems (3.3%) from our list, leaving us with 14,217 eukaryotic system candidates composed of 17,186 genes.

### Homology search of Mokosh, Eleos, Lamassu

We looked for homologs in eukaryotic genomes starting for the following primary HMM profiles: LmuA_effector_Protease, LmuB_SMC_Hydrolase_protease, MkoA, MkoB, LeoA, LeoBC (available at https://github.com/mdmparis/defense-finder-models). For each profile, the pipeline for homology search was similar. First, we searched the primary profile against our custom database of eukaryotic and prokaryotic proteins using hmmsearch (version 3.3.2), at an e-value threshold of 1e-3 (*32*). Prokaryotic hits were then marked as “paired” if they were found in proximity to their defense system partner(s), according to the rules of DefenseFinder, and as “isolated” otherwise. We selected a subset of these sequences, containing approximately 200 paired and 300 isolated prokaryotic hits, to perform downstream phylogenetic analyses. For both the paired and isolated groups, we picked half of the proteins among the top ranking hits to the reference hmm profile, and the other half by sampling uniformly along the list (Tables S3, S4, S5).

The subset of sequences was then filtered to remove duplicated proteins (using usearch v11.0.667 with the option -cluster_fast and -id 1) and aligned with mafft (version 7.505 with the option --auto) (*33*, *34*). The resulting alignment was trimmed with clipkit (version 1.3.0) using the kpic-gappy mode to keep informative and constant sites and removes sites with more than 90% of gaps (*35*). Phylogenetic trees were computed using IQTree (version 2.2.0.3) using model VT+F+I+G4 and 1000 ultrafast bootstraps (*36*), and visualized on the iTOL webserver (https://itol.embl.de). The trees were further annotated to display whether hits were paired or isolated, and whether the sequences were eukaryotic or prokaryotic. Clades were manually curated to include eukaryotic proteins distant from prokaryotic proteins with non-immune function (see fig. S1 and Table S3,S4,S5) . Their eukaryotic members were aligned using mafft. We performed a manual curation of the alignments by removing sequences that did not align well with the rest, recomputed the alignments, and manually truncated the N and C termini. These alignments were used to build secondary hmm profiles using hmmbuild (from hmmer version 3.3.2). All alignments and profiles are available at https://github.com/mdmparis/cury_mordret_et_al_supplementary_data.

Each of these secondary profiles was then used to search against the eukaryotic protein database using hmmsearch. We visualized the distribution of scores, and manually determined a score threshold to keep or discard these secondary hits (fig. S2). For each secondary profile, we composed a new subset of representative genes, composed of 300 isolated secondary hits, 100 paired eukaryotic secondary hits, all fusions (i.e. genes matched by the secondary hmm profiles of two different partners of the same system), all secondary hits in the human genome, and all the paired prokaryotic hits found in the previous search of the associated primary profile. We removed duplicates and performed the alignment in two steps: first, we aligned the prokaryotic sequences of the subset together using the procedure described above. The final alignment of bacterial proteins was performed using the option--maxiterate 1000 and --localpair (i.e. mafft-linsi). We then aligned the eukaryotic proteins to the bacterial alignment with maff-linsi parameters --add and --keeplength (*37*). We trimmed the alignment with clipkit, and generated trees as described above.

### Structural comparisons

Structures were either downloaded from the alphafold.ebi.ac.uk database, or computed using the ColabFold notebook (Table S7) (*38*–*40*). We created custom databases of structures and searched them using the easy-search command of foldseek version 3.915ef7d (*41*) (Table S7, https://github.com/mdmparis/cury_mordret_et_al_supplementary_data.) with default parameters. Structures were imported, realigned based on the Foldseek alignment, and visualized in the PyMol version 2.5.4 (*42*) (Schrödinger, LLC).

### Cell culture

Human embryonic kidney 293T cells (293T cells) and Vero cells were grown in Dulbecco’s modified Eagle’s medium (DMEM, Life Technologies) including 10% fetal calf serum and 100U/ml penicillin/streptomycin (Gibco). Prior to transfection, 293T cells were plated in medium without antibiotics.

### Viral stocks and plaque assay

Herpes Simplex virus 1 (HSV-1, KOS strain) was built by fusing VP26 fused to GFP (*43*). Sindbis virus was generated from the pTE32J infectious clone, as described elsewhere (*44*). Briefly, viral RNA was generated from *in vitro* transcription of linearized infectious clones, purified and electroporated in Vero cells. Viral stocks were amplified on 293T cells (HSV) or Vero cells (SINV) and titrated by plaque assay. Briefly, 200,000 Vero cells were seeded in 24-well plates and infected with 10-fold dilutions of sample for 1h at 37°C. Cells were then overlaid with DMEM containing 2% FCS, 100 U/ml penicillin-streptomycin and 0.8% agarose. After 3 days (SINV) or 4 days (HSV-1), cells were fixed with 10% formalin and visualized by crystal violet staining.

### Plasmids, transfection and viral infections

Human proteins of interest were synthetized through Genscript and integrated in a pcDNA3.1(+) backbone. GIMAP5 and 6 were tagged with a flag tag and a HA tag at the N-terminus, respectively, whilst EFHD2 was equipped with a flag tag (N-terminus), CTRC with a HA tag (C-terminus) and FHAD1 with a V5 tag (C-terminus). 100 ng of each plasmid was transfected in 293T cells using Lipofectamine 2000 (Thermo Fisher Scientific) following the manufacturer’s instruction. One day after plasmid transfection, cells were infected with HSV-GFP or SINV-GFP at multiplicity of infection 0.5. Viral supernatant was collected at the indicated time points for viral particle titration, or cells were collected at 48h post-infection for flow cytometry analysis.

### Flow cytometry

Cells were collected in a V-shape bottom 96-well plate (Corning), and resuspended in 50 ul PBS supplemented with DAPI. FIX & PERM kit (Thermo Fisher Scientific, GAS004) was used according to the manufacturer’s instructions. Proteins of interest were visualized using an anti-Flag-PE (BioLegend, 637310), an anti-HA-Alexa647 (BioLegend, 682404) and an anti-Flag-PE-Cy7 (Thermo Fisher Scientific, 25-6796-42) at a 1:300 dilution. Analysis were performed on a FACSVerse Cell Analyser (BD Biosciences) and results were analyzed with FlowJo (TreeStar).

### Data visualization

Trees were drawn with ITOL (https://itol.embl.de). Schematics of Eukaryotic organisms were taken from https://beta.phylopic.org/. Graphs were plotted using matplotlib (https://matplotlib.org) and seaborn (http://seaborn.pydata.org/index.html). Web logos of alignment were built using webserver of weblogo (*45*).

## Acknowledgments

We thank Nicolas Manel and Xavier Lahaye for the kind gift of herpes simplex virus. The bioinformatics analysis were performed on the Core Cluster of the Institut Français de Bioinformatique (IFB) (ANR-11-INBS-0013). We are grateful to members of the MDM lab, U1284, U932 as well as mentors and friends for their useful comments on earlier version of the manuscript.

## Author contributions

All the authors designed experiments and analyzed data. J.C., E.M. and F.T. performed data curation and formal analysis. V.H.T performed experiments. J.C., E.M., G.O., E.Z.P. and A.B. wrote the manuscript. A.B. and E.Z.P. acquired funding.

## Data and materials availability

All the data, including phylogenetic trees, HMM profiles and protein structures generated with AlphaFold, are available at https://github.com/mdmparis/cury_mordret_et_al_supplementary_data.

