## Supplementary Materials for "Conservation of antiviral systems across domains of life reveals novel immune mechanisms in humans"

Cury, Mordret *et al.*

### Supplementary Materials

**Supplementary Materials for this manuscript include the following:**

Supplementary Table 1: DefenseFinder hits
Supplementary Table 2: Summary of homology analysis
Supplementary Table 3: Hits for the Mokosh system (**a.** prokaryotic and eukaryotic hits from search using prokaryotic profiles. **b.** eukaryotic hits from search using eukaryotic profiles)
Supplementary Table 4: Hits for the Eleos system (**a.** prokaryotic and eukarotic hits from search using prokaryotic profiles. **b.** eukaryotic hits from search using eukaryotic profiles)
Supplementary Table 5: Hits for the Lamassu system (**a.** prokaryotic and eukarotic hits from search using prokaryotic profiles. **b.** eukaryotic hits from search using eukaryotic profiles)
Supplementary Table 6: Human homologs of Mokosh, Eleos and Lamassu
Supplementary Table 7: TM-scores of structural comparisons

All the data, including phylogenetic trees, alignements, HMM profiles, protein structures are available for download at the following link:
https://github.com/mdmparis/cury_mordret_et_al_supplementary_data

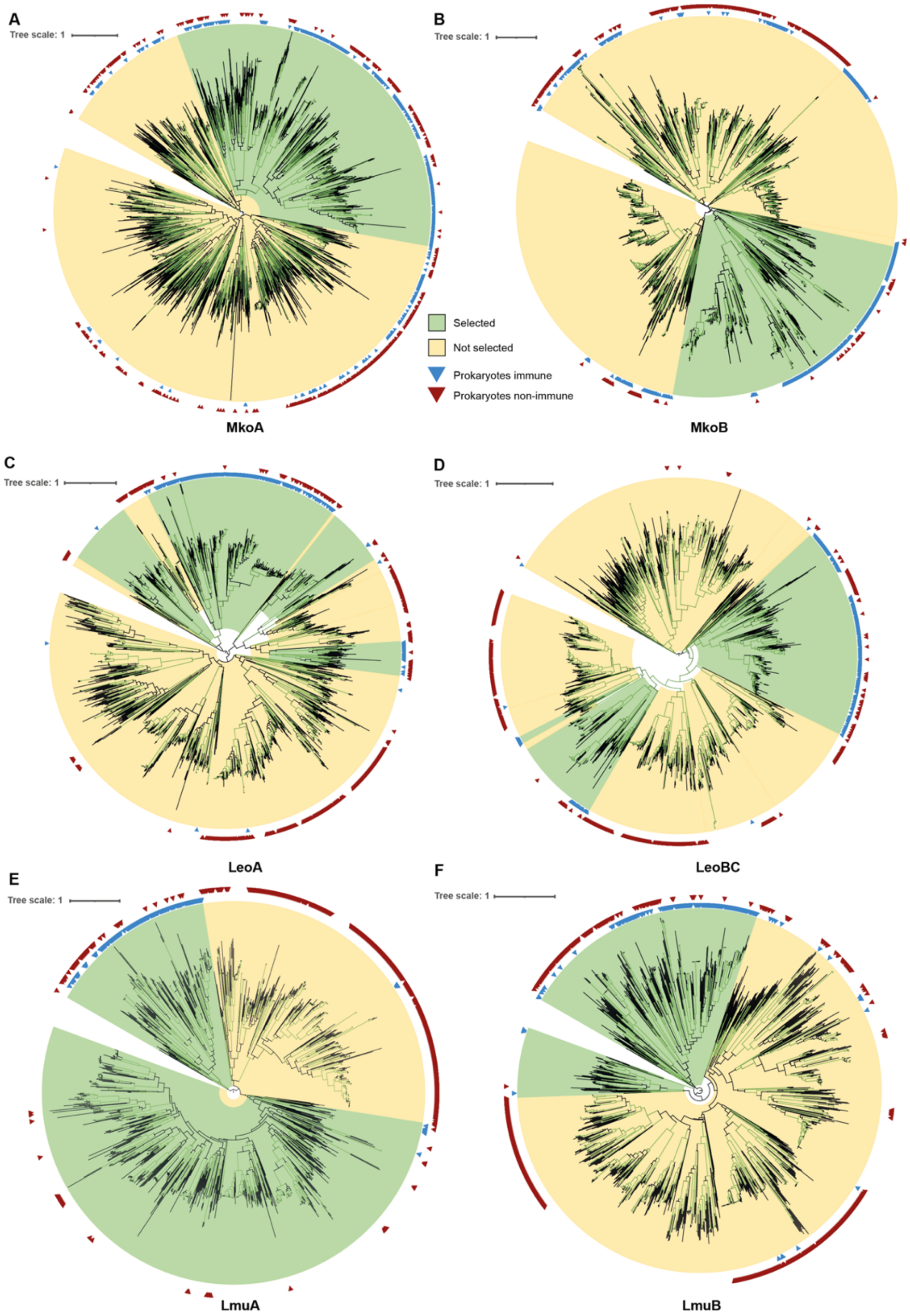


**Fig. S1. Phylogenetic trees used to select proteins destined to build eukaryotic HMM profiles.** Green branches correspond to ultrafastboostrap >95 (in IQTree). Interior ring indicates prokaryotic hits with colocalized antiphage partner (blue). Exterior ring displays prokaryotic hits without colocalized partner (purple, hits not detected by DefenseFinder). Clades selected to build the HMM eukaryotic profile are indicated in green. Some green clades contain only Bacteria. Clades in yellow are removed from HMM profile construction.

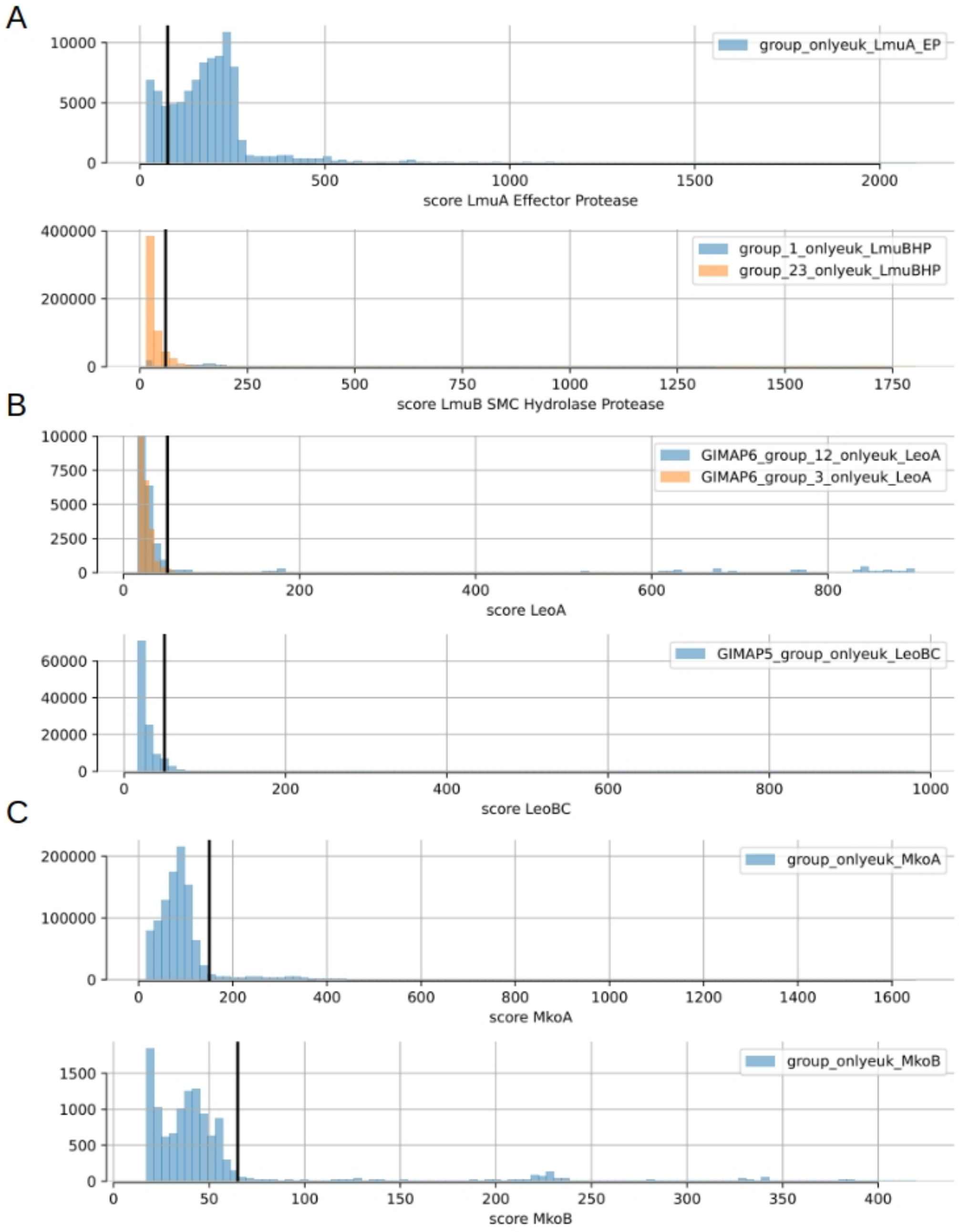


**Fig. S2. Score distribution used to select hits based on eukaryotic HMM profiles.**
Positive hits obtained from the analysis using new eukaryotic profiles were selected with the indicated threshold (vertical bar), determined by the distribution of scores. When two different clades were selected, two profiles were used (represented in blue and orange, see fig. S1). Labels int the top right corner correspond to the name of the eukaryotic HMM profiles.

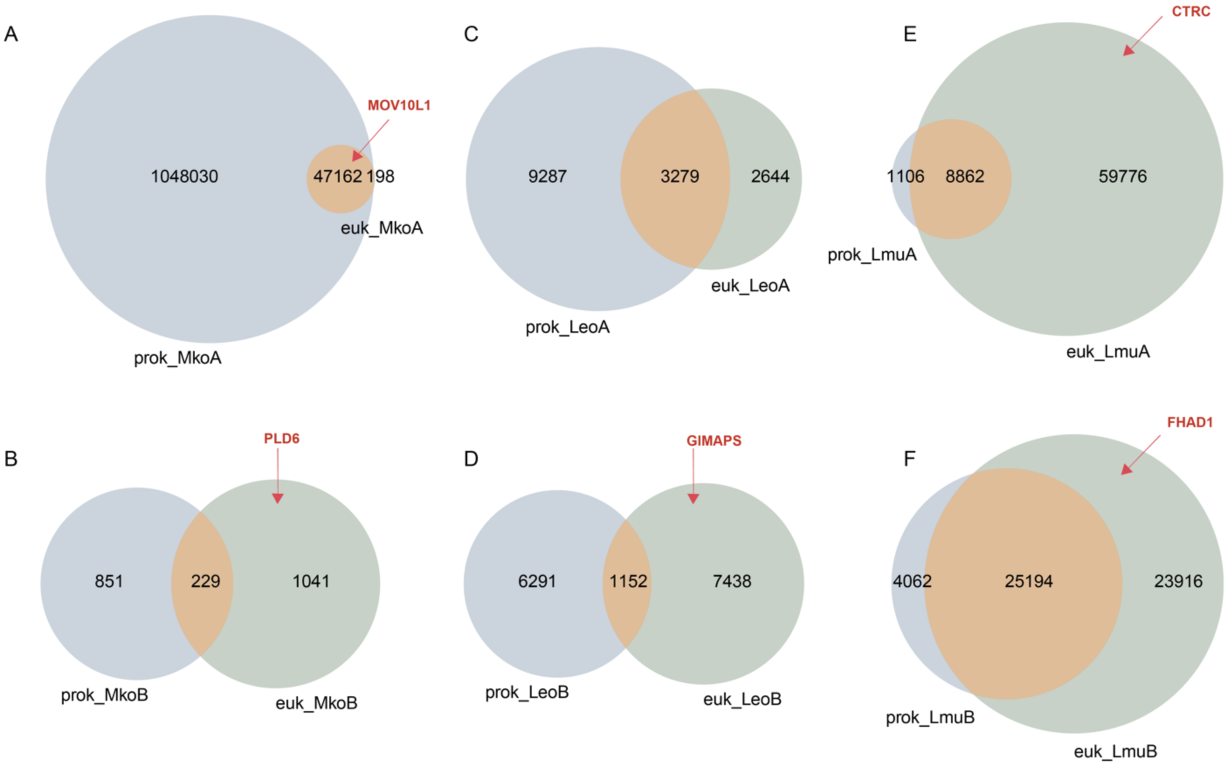


**Fig. S3. Hits obtained from homology analysis using prokaryotic or eukaryotic HMM profiles.** Venn diagrams represent hits from both analyses. In blue, hits starting from prokaryotic HMM profiles, in green hits starting from eukaryotic HMM profiles, in orange hits found by both analysis. Human genes explored in the study are indicated with a red arrow (all detected using eukaryotic HMM only except for MOV10L1, isolated in both analysis).

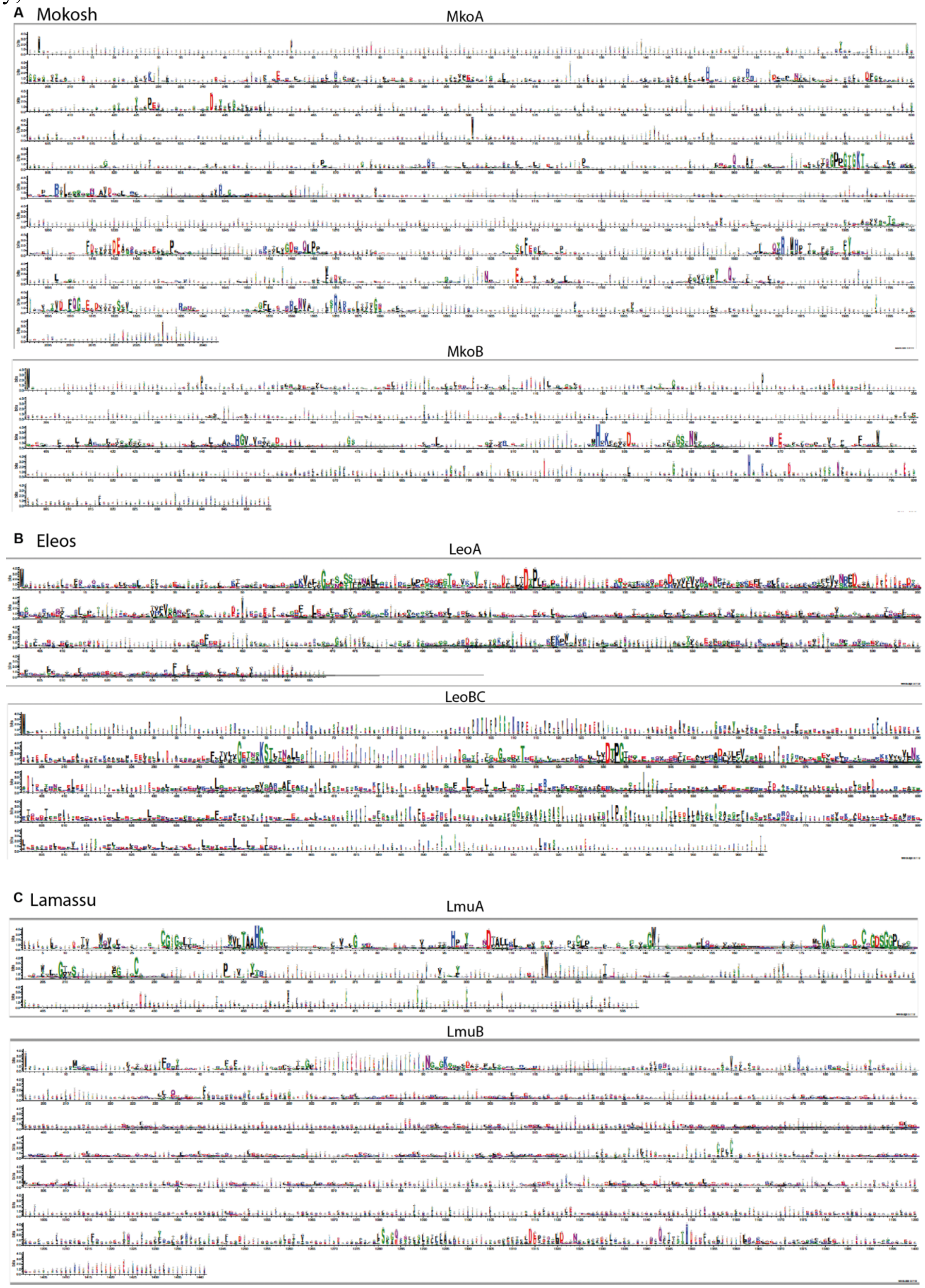


**Fig. S4. Weblogos of trimmed alignments of final hits.**
Prokaryotic hits were detected using DefenseFinder. Eukaryotic hits were detected using eukaryotic HMM profiles described in the text and available on the GitHub of the supplementary data.

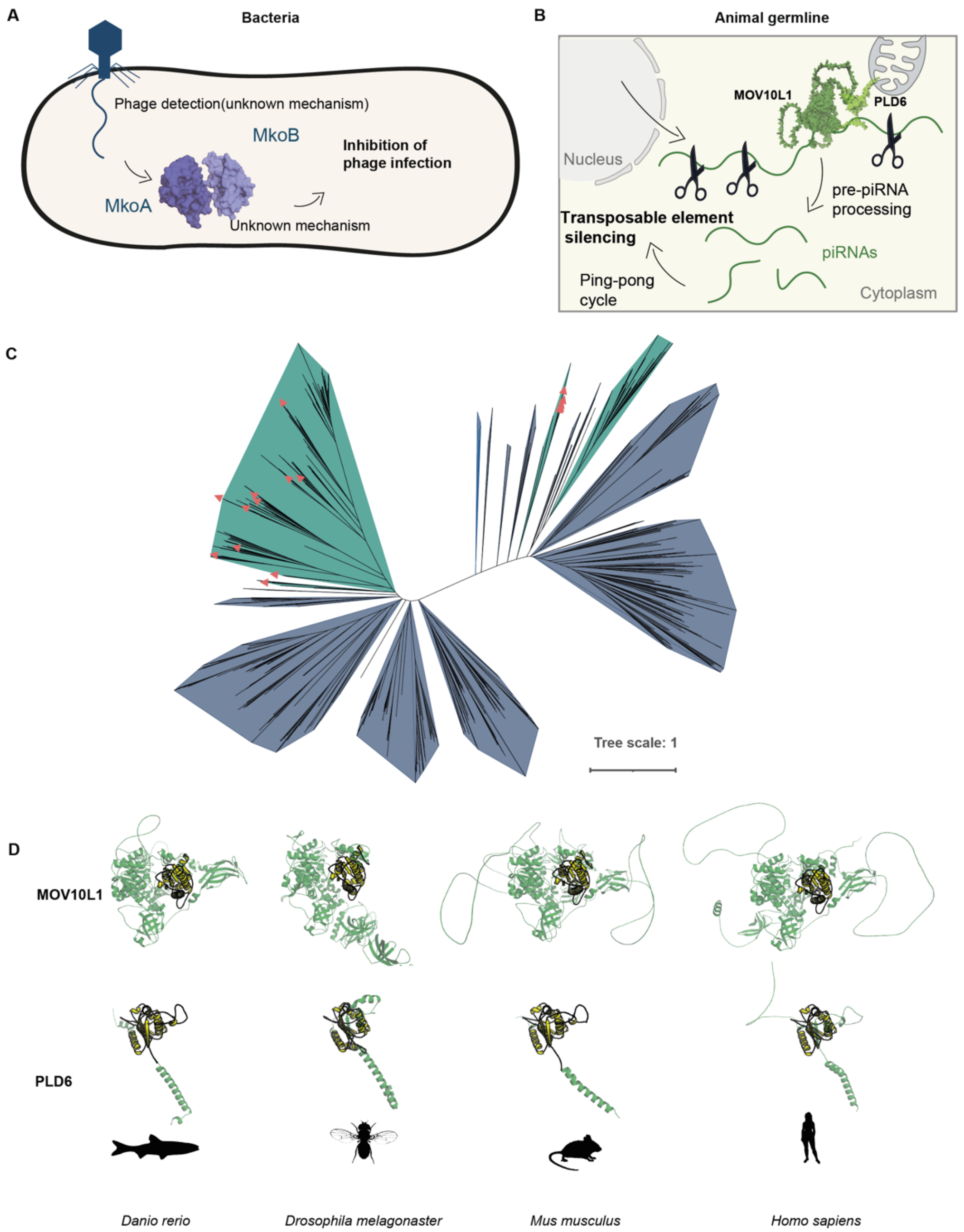


**Fig. S5. Homology between antiphage Mokosh and the animal piRNA pathway.**
(**A**) In bacteria, the Mokosh system detects and thwarts phage infection through unknown mechanisms. (**B**) In the animal germline, repression of deleterious transposon expression relies on the cleavage of precursor piRNAs (pre-piRNAs) by MOV10L1 and PLD6 to generate piRNAs that are used to sequence-specifically target transposons. (**C**) MkoA phylogenetic tree with its eukaryotic hits. Human hits are indicated with red triangles. (**D**) Structural comparisons of MOV10L1 (top) and PLD6 (bottom) of various eukaryotic species (in green) with the yjhR protein. For MOV10L1 (respectively PLD6) homologs, the yellow domain corresponds to the protein's optimal alignment to the MkoA (respectively MkoB) domain of yjhR. Structures were predicted using AlphaFold. PDB identifiers and TM-scores can be found in Table S7.

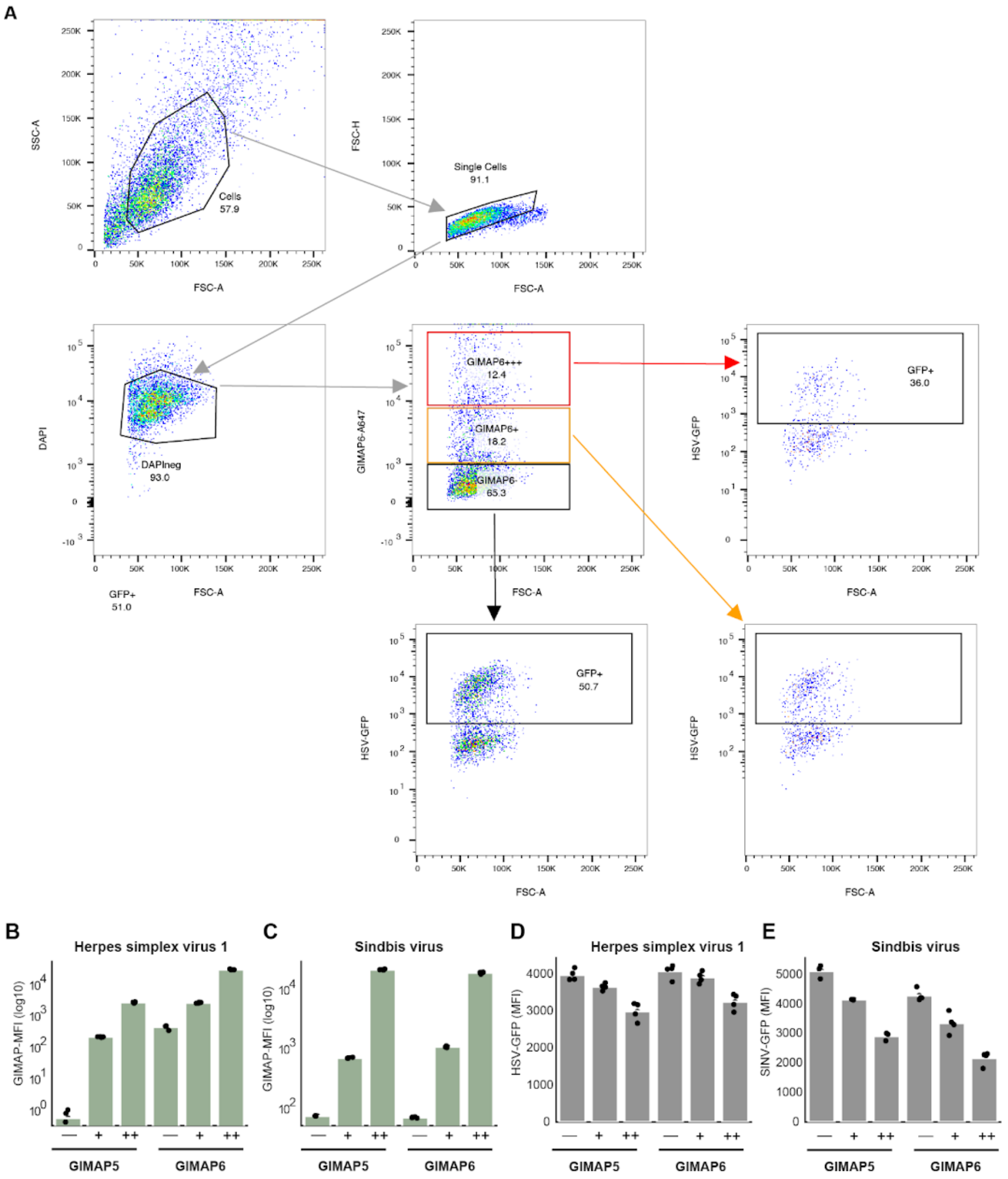


**Fig. S6. Antiviral activity of human GIMAPs.**
(**A**) Flow cytometry analysis of 293T cells transfected with a plasmid encoding for GIMAP6 tagged with HA and infected with HSV-1 coding for GFP. (**B-E**) 293T cells were transfected with a plasmid encoding for GIMAP5 or 6, and infected with HSV-1 (B,D) or SINV (C,E). GIMAP expression is plotted as mean fluorescent intensity (B,C). GFP mean fluorescence intensity is measured as a proxy of HSV-1 (D) or SINV (E) replication.

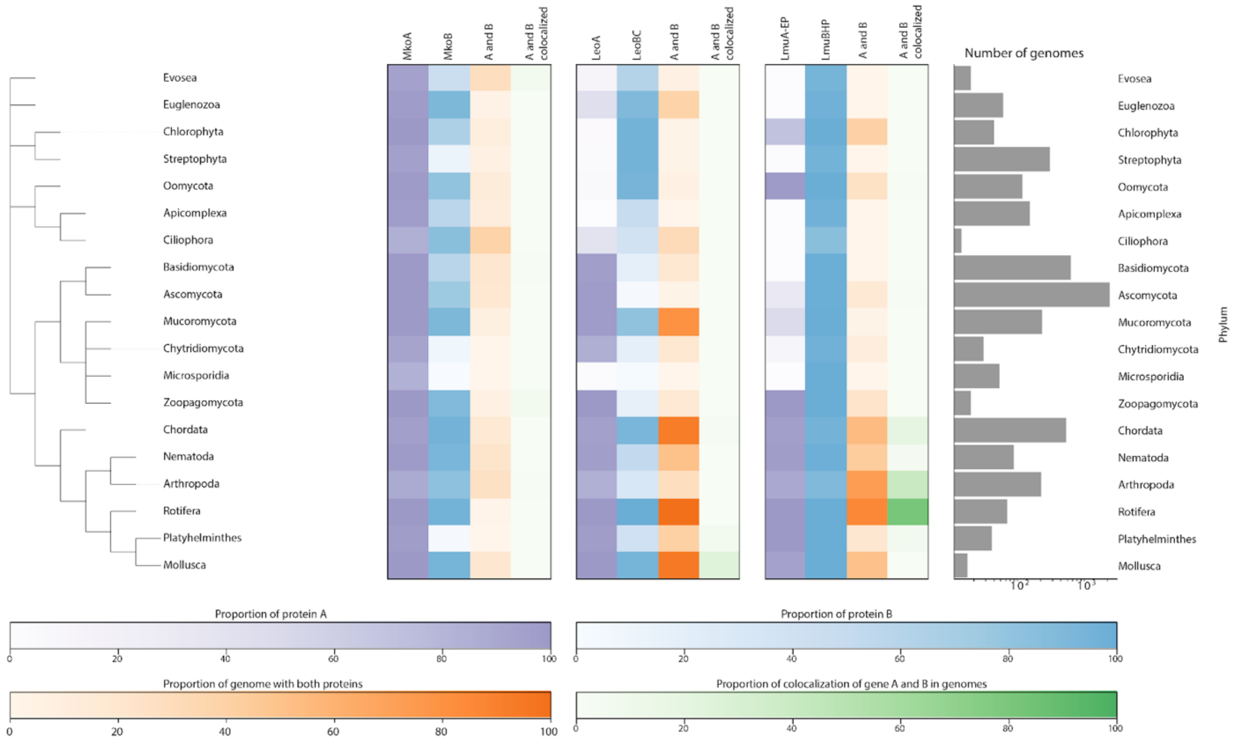


**Fig. S7. Distribution of Mokosh, Eleos and Lamassu in eukaryotic phyla.**
Heatmaps represent the proportion of genomes with at least one protein showing homology to antiphage proteins. For a given system, protein A is colored in purple and protein B in blue. Third colum (“A and B”) represent the proportion of genomes of a given phyla in which protein A and B are concomitantly detected. Fourth column (“A and B colocalized”) quantifies the proportion of genomes showing colocalization for genes coding for proteins A and B. Graph on the right represents number of genomes in each phyla (log scale). Selected phyla display more than 10 genomes in the eukaryotic genomic database.

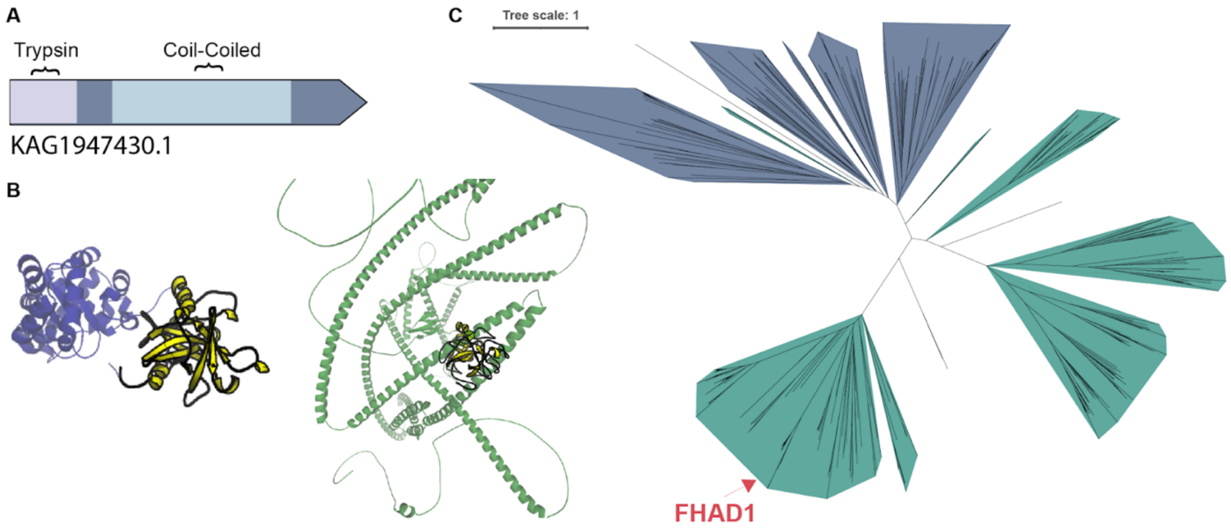


**Fig. S8. Conservation of antiphage Lamassu in eukaryotes, including humans.**

(**A**) Schematic of a fusion protein detected in *Pimephales promelas* (fathead minnow, a fish from the *Cyprinidae* family), bearing a trypsin domain (homolog of LmuA) and a coil-coiled domain (homolog of LmuB). (**B**) Computed structure of a Lamassu fusion from *E. coli* (blue) and its homology with the trypsin domain of *Pimephales promelas* fusion protein (green). Optimal local alignment between the two structures (as determined by foldseek) is represented in yellow. TM-score = 0.67. (**C**) Phylogenetic analysis combining prokaryotic LmuB with its eukaryotic homologues (see Material and Methods). Branches are colored according to the kingdom (blue for prokaryotes, green for eukaryotes). Human homolog FHAD1 is indicated in red.

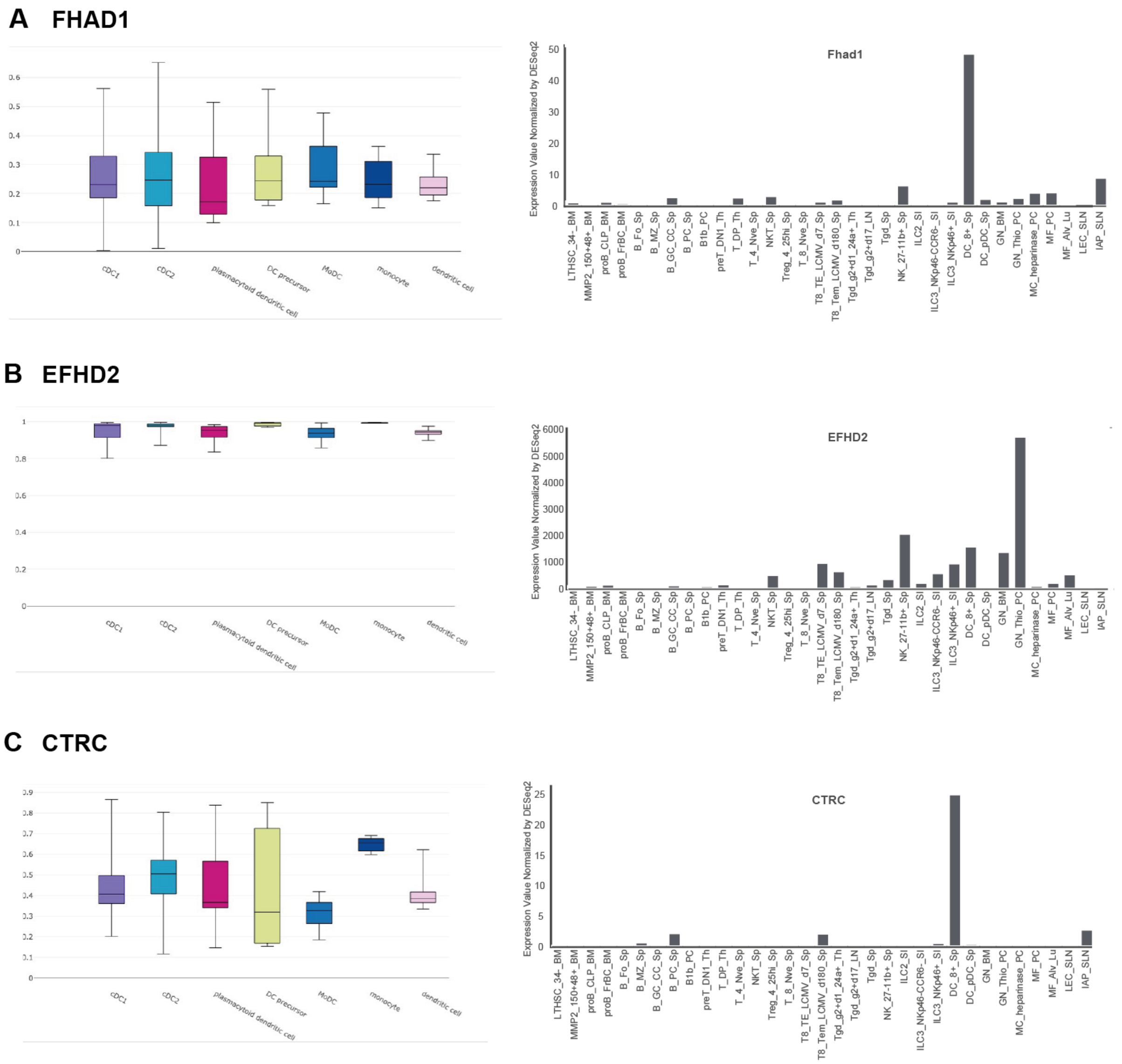


**Fig. S9. Expression of FHAD1, EFHD2 and CTRC in immune cells.**
(**A-C**) Transcript expression for FHAD1 (A), EFHD2 (B) and CTRC (C) extracted from an atlas of myeloid and dendritic cell subsets (left panels, (*34*)) and from the ImmGen database (right panels, http://rstats.immgen.org/Skyline/skyline.html).

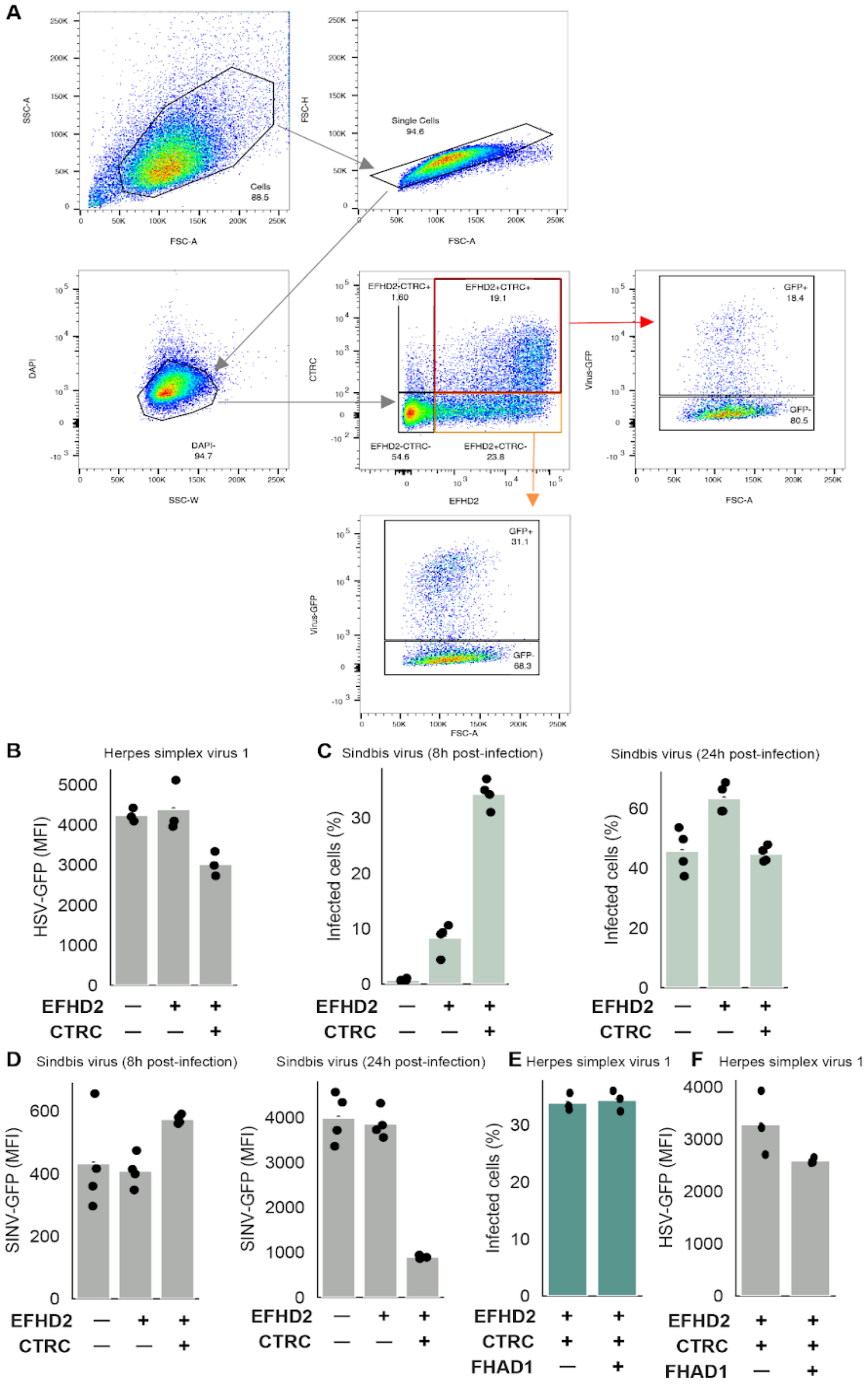
A
SSC-A
FSC-A
Cells
88.5
FSC-H
Single Cells
94.6
DAPI
SSC-W
DAPI-
94.7
CTRC
EFHD2
EFHD2-CTRC+
1.60
EFHD2+CTRC+
19.1
EFHD2-CTRC-
54.6
EFHD2+CTRC-
23.8
Virus-GFP
GFP+
18.4
GFP-
80.5
GFP+
31.1
GFP-
68.3
B
Herpes simplex virus 1
HSV-GFP (MFI)
EFHD2
CTRC
C
Sindbis virus (8h post-infection)
Infected cells (%)
Sindbis virus (24h post-infection)
D
Sindbis virus (8h post-infection)
SINV-GFP (MFI)
Sindbis virus (24h post-infection)
E
Herpes simplex virus 1
FHAD1
F
Herpes simplex virus 1

**Fig. S10. Antiviral activity of human EFHD2 and CTRC.**
(**A, B**) Flow cytometry analysis of 293T cells transfected with a plasmid encoding for EFHD2 tagged with Flag, CTRC tagged with HA and infected with HSV-1 coding for GFP. Mean fluorescence intensity (MFI) of GFP is plotted in (B). (**C,D**) 293T cells transfected with plasmids encoding EFHD2 and CTRC were infected with SINV-GFP. Percentage of infected cells, as visualized by GFP-positive cells (C), as well as GFP MFI, indicative of viral replication (D), was measured by flow cytometry at 8 and 24 hours post infection. (**E,F**) 293T cells transfected with plasmids encoding FHAD1, EFHD2 and CTRC were infected with HSV-1 encoding for GFP. Percentage of infected cells (E), as well as GFP MFI (F), was monitored by flow cytometry at 48 hours post infection.
